# The Mlh1-Pms1 endonuclease uses ATP to preserve DNA discontinuities as strand discrimination signals to facilitate mismatch repair

**DOI:** 10.1101/2024.06.13.598860

**Authors:** Jonathan M. Piscitelli, Scott J. Witte, Yasmine S. Sakinejad, Carol M. Manhart

**Affiliations:** Department of Chemistry, Temple University, Philadelphia, Pennsylvania, 19122, USA

**Author notes:** Joint Authors (Jonathan M. Piscitelli, Scott J. Witte).

## Abstract

In eukaryotic post-replicative mismatch repair, MutS homologs (MSH) detect mismatches and recruit MLH complexes to nick the newly replicated DNA strand upon activation by the replication processivity clamp, PCNA. This incision enables mismatch removal and DNA repair. Biasing MLH endonuclease activity to the newly replicated DNA strand is crucial for repair. In reconstituted *in vitro* assays, PCNA is loaded at pre-existing discontinuities and orients the major MLH endonuclease Mlh1-Pms1/MLH1-PMS2 (yeast/human) to nick the discontinuous strand. *In vivo,* newly replicated DNA transiently contains discontinuities which are critical for efficient mismatch repair. How these discontinuities are preserved as strand discrimination signals during the window of time where mismatch repair occurs is unknown. Here, we demonstrate that yeast Mlh1-Pms1 uses ATP binding to recognize DNA discontinuities. This complex does not efficiently interact with PCNA, which partially suppresses ATPase activity, and prevents dissociation from the discontinuity. These data suggest that in addition to initiating mismatch repair by nicking newly replicated DNA, Mlh1-Pms1 protects strand discrimination signals, aiding in maintaining its own strand discrimination signposts. Our findings also highlight the significance of Mlh1-Pms1’s ATPase activity for inducing DNA dissociation, as mutant proteins deficient in this function become immobilized on DNA post-incision, explaining *in vivo* phenotypes.

**GRAPHICAL ABSTRACT:** 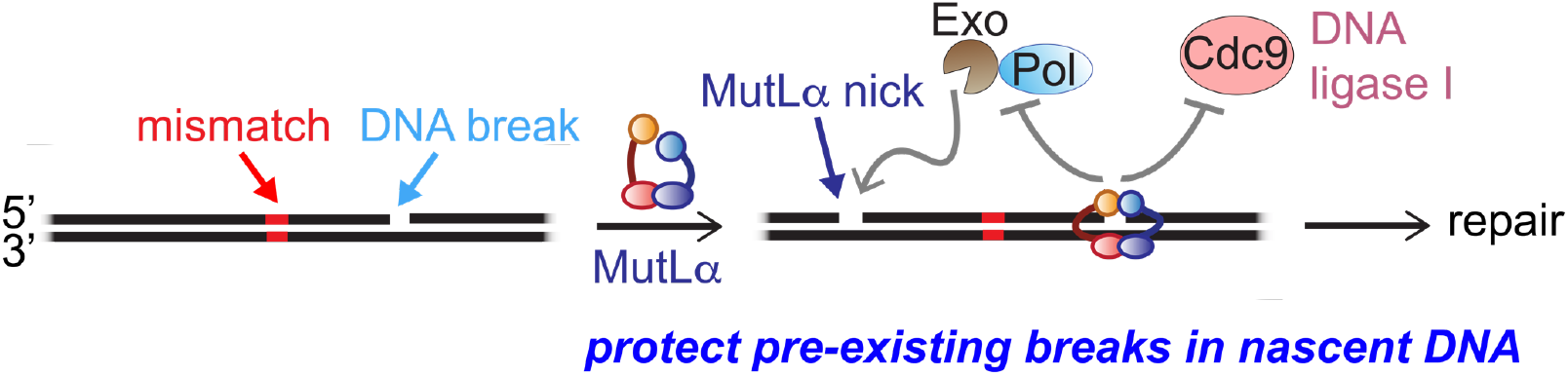

## INTRODUCTION

DNA mismatch repair is a highly conserved process that improves the overall fidelity of DNA replication by removing bases misincorporated by DNA polymerases in the nascent DNA strand (1). Consistent with this, defects in mismatch repair proteins are associated with increased mutation rates and, in humans, increased incidence of cancer (1, 2).

In eukaryotes, the major mismatch repair pathway involves recognition of mismatches by either the Msh2-Msh6 or Msh2-Msh3 complex. These proteins then recruit the heterodimeric endonuclease Mlh1-Pms1 (in the yeast nomenclature, MLH1-PMS2 in mouse and human) which nicks the nascent DNA strand upon activation through interactions with the proliferating cell nuclear antigen (PCNA) protein. Mlh1-Pms1’s nick is critical for mismatch repair and several, overlapping pathways have been suggested for how this nick is used to remove the mismatch (1, 3–6). For exonuclease-dependent mismatch repair, the 5’-terminus of the Mlh1-Pms1 nick is used by Exo1 to begin excision and remove a segment of DNA which includes the mismatch. The resultant gap is then repaired by RPA, Polδ, and a ligase. For exonuclease-independent repair, the 3’-end of the Mlh1-Pms1-generated nick is used as an initiation point for Polδ to synthesize DNA towards the mismatch, displacing a segment of the DNA strand that contains the mismatch as a flap, which is cleaved by the Rad27 flap endonuclease (7). Another pathway has been suggested where Mlh1-Pms1 can iteratively nick the nascent strand near the mismatch. In this scenario, the mismatch is removed either through diffusion of the oligonucleotides formed by the Mlh1-Pms1 repetitive nicking, by Exo1 exonuclease activity, or strand displacement synthesis by Polδ (6, 8, 9).

These pathways all rely on Mlh1-Pms1 to generate a nick in the newly replicated DNA strand and not in the template strand. The mechanistic underpinnings for how this bias is achieved are unclear and have been highly sought-after largely through biochemical assays. Partially and fully reconstituted mismatch repair assays require plasmid-based substrates to both mimic chromosomal DNA where most of the genome is not in the vicinity of a DNA end, and to account for observations that MLH endonucleases have high affinity for and require large DNA substrates to be endonuclease active (6, 10–17). In partially reconstituted mismatch repair assays dependent on the Mlh1-Pms1/PMS2 endonuclease activity, this activity is shown to have strand bias only when there is a pre-existing nick in the model plasmid substrate (14, 15). In minimal systems using *Saccharomyces cerevisiae* and human proteins, it was shown that Mlh1-Pms1’s bias towards nicking the strand with the pre-existing discontinuity was dependent on PCNA and its loading protein replication factor C (RFC) (14, 15, 17). These data suggest that the pre-existing discontinuities in these assays serve as sites for PCNA loading and the orientation of PCNA relative to this discontinuity directs Mlh1-Pms1/PMS2 endonuclease activity.

A direct role for PCNA in dictating strand bias is also suggested in experiments using *Xenopus* egg extracts. In this system, PCNA was loaded by RFC onto a plasmid DNA with a discontinuity which was then ligated closed, and the PCNA-bound plasmid was then used as a substrate for *ex vivo* mismatch repair (18). In these assays, mismatch repair was biased towards the strand that previously contained the discontinuity suggesting that PCNA orientation is sufficient for dictating strand bias, but the discontinuity itself is not required.

Despite PCNA’s role in activating and directing repair *in vitro*, microscopy experiments in *S. cerevisiae* show that Mlh1-Pms1 foci are distant from replication centers during repair *in vivo* (19). It is unclear whether PCNA used in mismatch repair is a remnant of the replication fork or is loaded onto DNA by RFC during repair. If the latter, strand discontinuities on a newly replicated strand could act as PCNA loading sites. Regardless of what mechanistically pre-existing nicks are used for in DNA mismatch repair, both the leading and lagging strands can contain discontinuities (1, 20, 21) and pre-existing nicks have been established *in vivo* as being important for conveying strand bias. This was shown in experiments where the Cdc9 ligase was overexpressed and was antagonistic to mismatch repair in *S. cerevisiae* (22). These data suggest that prematurely removing discontinuities from newly replicated DNA eliminates the temporal window between DNA replication and mismatch repair, increasing mutation rates. It therefore follows, that these discontinuities need to be preserved for the duration of time between DNA replication until post-replicative mismatch repair occurs to ensure the fidelity of the pathway. How DNA discontinuities are protected to promote strand discrimination and efficient mismatch repair has not been explored, but our data using purified Mlh1-Pms1 from *S. cerevisiae* suggest that Mlh1-Pms1 plays a direct role in maintaining DNA discontinuities as strand discrimination signals.

The Mlh1-Pms1 heterodimer has several notable biochemical properties that it uses to facilitate DNA mismatch repair. The homologous Mlh1 and Pms1 subunits contain conserved globular amino terminal domains and globular carboxy-terminal domains which are connected via poorly conserved intrinsically disordered linker regions (23). The Mlh1 and Pms1 subunits primarily dimerize through their carboxy-terminal domains and this dimerization interface overlaps with the endonuclease site, which is primarily located in the carboxy-terminal domain of the Pms1 subunit (24). Mlh1-Pms1’s endonuclease activity has been shown previously to be activated mainly via interactions between the carboxy-terminal domain of Pms1 and PCNA (16). The globular amino-terminal domains of each subunit contain conserved ATPase sites (25–27). Although, ATP has been shown to not be absolutely required for endonuclease activity *in vitro*, it has been shown to have a stimulatory effect on this activity (13, 28) and to promote transient dimerization between the Mlh1 and Pms1 subunits via the amino-terminal domains (25, 26, 29).

Previous research on the ATPase activities of *S. cerevisiae* Mlh1-Pms1 suggests that disrupting the site in either subunit can elevate mutation rates and induce a null-like phenotype *in vivo* (25, 27). Atomic force microscopy experiments and partial proteolysis assays using purified Mlh1-Pms1 have revealed significant conformational changes induced by ATP binding and hydrolysis via the intrinsically disordered regions (27, 30). *In vitro* biochemical assays using yeast and *E. coli* homologs suggest that ATP hydrolysis regulates the interactions between the protein and DNA (28). These findings are consistent with assays involving yeast, human, and bacterial homologs, which have revealed that Mlh1-Pms1’s ATPase activity is stimulated by DNA (16, 28, 29), and further enhanced by interactions with PCNA (16, 28). Although not directly observed, the proposed large conformational changes in response to ATP binding and hydrolysis are believed to facilitate a recycling mechanism wherein Mlh1-Pms1 cleaves DNA in mismatch repair, dissociates during ATP hydrolysis, and subsequently undergoes rebinding and reactivation to iteratively nick DNA. However, the mechanisms by which these signals are transmitted within the protein and how ATPase deficiencies lead to failures in this process, resulting in null-like phenotypes, remain unclear as iterative nicking does not seem to be required for efficient mismatch repair (6, 8, 9, 28). Due to this, the role of Mlh1-Pms1’s ATPase activity and why it is critical for mismatch repair is unclear.

In this study, we determined that Mlh1-Pms1 uses ATP-binding to protect DNA discontinuities. Upon discontinuity recognition, the protein loses its ability to efficiently interact with PCNA which in turn inhibits Mlh1-Pms1’s endonuclease activity and attenuates its ATPase activity and ability to dissociate from DNA. This feature can be used by Mlh1-Pms1 to protect discontinuities as strand discrimination signals and to prevent inappropriate excision and replication initiating at single strand breaks that may be present on nascent DNA due to Okazaki fragment synthesis and non-processive DNA replication in addition to preventing Mlh1-Pms1 from making additional nicks in the DNA when mismatch repair is not signaled. Furthermore, our investigation suggests a plausible explanation for the observed *in vivo* defects in ATPase mutants—although these mutants retain the ability to nick DNA, they become trapped at the nick site and are unable to dissociate. Consequently, this impairs access to the nick by downstream factors required for mismatch removal.

## MATERIAL AND METHODS

### Proteins and DNA Substrates Used in This Study

Wild-type yeast Mlh1-Pms1 was expressed and purified as described previously (31). Mlh1-Pms1 ATPase mutations (*mlh1N35A-pms1N34A, mlh1N35A-Pms1*, and *Mlh1-pms1N34A*) were generated by Q5 mutagenesis using primers CMO175 and 176 for the mutation in Mlh1 using plasmid pMH1 and primers CMO177 and 178 for the mutation in Pms1 using plasmid pMH8 (Table S1). Plasmids pMH1 and pMH8 that express yeast Mlh1 and Pms1, respectively, were a gift from Thomas Kunkel’s lab. Yeast PCNA and RFC were expressed and purified according to previous reports (32, 33).

Unless otherwise noted 2.7 kb closed circular substrates is pUC18 and the 4.3 kb closed circular substrate is pBR322 (Invitrogen). Linear plasmid-based substrates were generated by digestion with HindIII restriction enzyme (NEB) according to the manufacturer’s instructions followed by heat inactivation. Substrates were then isolated by P-30 spin-column (Bio-Rad). Plasmid-based substrates with nicks were generated by incubating with nicking restriction endonucleases Nb.BspQI (1 nick), Nb.BsrDI (2 nicks), Nb.BtsI (3 nicks) Nt.BstNBI (4 nicks) and Nb.BsrDI was combined with Nb.BtsI (5 nicks) (Figure S1) according to the manufacturer’s instructions followed by spin-column purification. Circularized DNA without nicks was generated incubating 5 units of Topoisomerase I with the DNA for 24 hours and deactivated according to the manufacturer’s instructions.

For gel shift assays on oligonucleotides, CMO169 was radiolabeled with [γ-^32^P]-ATP (Perkin Elmer) by T4 polynucleotide kinase (NEB) (Table S1). Unincorporated nucleotide was removed using G-25 spin-columns (Cytiva) per the manufacturer’s instructions. Radiolabeled CMO169 was then annealed with either CMO168 to create the intact duplex substrate or with CMO170 and CMO184 to create a single strand break substrate in 10 mM Tris-HCl pH 7.5, 0.5 M NaCl, and 10 mM EDTA, by incubating at 99°C for 5 minutes and slowly cooling to room temperature over the span of 3 hours (Table S1).

### Electrophoresis Mobility Shift Assays

DNA substrates (3.8 nM final concentration) were combined in 20 μL reactions on ice with varying concentrations of wild-type or mutant Mlh1-Pms1 in Buffer A, which contains 20 mM HEPES-KOH, 0.1 mM DTT, 2 mM MgCl_2_, 40 μg/mL BSA, and 6% glycerol (final concentrations). Where indicated, 500nM of PCNA or 0.5 mM ATP was also included. Reactions were incubated for 10 minutes at room temperature and loaded into a 1% (w/v) agarose gel in a buffer containing 40 mM Tris, 20 mM acetate, and 1 mM EDTA (1X TAE) buffer that was prechilled for 1 hour at 4°C. The gel was resolved on ice for 2 hours at 45 volts and stained in 300 mL of 1X TAE containing a final concentration of 10 μg/mL ethidium bromide. For assays using radiolabeled oligonucleotide duplexes, reactions were assembled in Buffer A with the addition of 120 mM NaCl. Where indicated, 0.5 mM ATP was also included. After a 10 minute incubation at 4°C, products were resolved by 4% polyacrylamide gel run in 1X TBE (89 mM Tris, 89 mM boric acid, 2 mM EDTA) at 110 V for 1 hour at 4°C. Gels were visualized by phosphorimaging.

### ATPase Assays

ATPase assays were performed as previously described and analyzed by thin layer chromatography (TLC) (16, 26, 27). Briefly, 10 μL reactions were combined containing 400 nM Mlh1-Pms1, 3.8 nM pBR322 DNA substrate, 500 nM PCNA where indicated, and 100 μM and [γ-^32^P]-ATP (Perkin Elmer) in a buffer containing 20 mM HEPES-KOH pH 7.5, 2.5 mM MnSO_4_, 2.0 mM MgCl_2_, 20 mM KCl, 1% glycerol, 0.2 mg/mL BSA (final concentrations). Reactions were incubated for 45 minutes at 37°C. After incubating, 1 μL of reaction was spotted onto a polyethylenimine cellulose plate (Sigma-Aldrich). Plates were developed for 1 hour using a solution of 1 M formic acid and 0.8 M LiCl_2_ and visualized by phosphorimaging.

### Endonuclease Assays

Endonuclease reactions were performed as previously described (12, 28, 34). Briefly, 20 μL reactions were combined containing varying concentrations of either wildtype or mutant Mlh1-Pms1, with 3.8 nM DNA substrate, in a buffer containing 20 mM HEPES-KOH, 20 mM KCl, 1% glycerol, 0.2 mg/mL BSA, 2.5 mM MnSO_4_ and 0.50 mM ATP (final concentrations). Reactions were incubated at 37°C for 60 min and stopped using stop mix (final concentrations: 1% SDS, 14 mM EDTA, 0.96 units proteinase K). Reactions were run on a 1.2% (w/v) agarose gel in 1X TAE for 1 hour at 100 V. For endonuclease assays on linear DNA substrates, reactions were denatured with 30 mM NaOH, 1 mM EDTA, 3% glycerol, and 0.02% bromophenol blue (final concentrations), then heated to 70°C for 5 minutes, and placed on ice for 3 minutes. Endonuclease products were then resolved on a 1% (w/v) agarose gel containing 30 mM NaCl and 2 mM EDTA, in a solution of 30 mM NaOH and 2 mM EDTA at 50 V for 2.5 to 3 hours. Denaturing gels were neutralized with 0.5 M Tris-HCl (pH 7.5) for 30 minutes. All gels were stained, visualized, and quantified as described for electrophoretic mobility shift assays.

### Exonuclease Inhibition Assay

For exonuclease protection assays in Figure 5A-C, 3.8 nM DNA was combined with varying concentrations of wild-type or mutant Mlh1-Pms1, along with 100 nM RFC, and 500 nM PCNA in a buffer containing 20 mM HEPES-KOH, 20 mM KCl, 1% glycerol, 0.2 mg/mL BSA, 2.5 mM MnSO_4_ and 0.25 mM ATP. The reaction was incubated at 37°C for 1 hour to promote endonuclease activity. Following this incubation, 8 units of T7 exonuclease (NEB) were added. Reactions were incubated at 37°C for an additional 30 min and stopped using stop mix (final concentrations: 1% SDS, 14 mM EDTA, 0.96 units proteinase K) as described above. Reactions were then incubated at 70°C for 30 min to stop the exonuclease reaction. Reactions were then analyzed by 1% (w/v) agarose gel run at 100 V for 60 min, stained, visualized, and quantified as described above. For exonuclease protection assays in Figure 5F, 3.8 nM of 2.7 kb plasmid DNA containing 1 nick (generated with Nt.BspQI) was incubated with 0 nM, 50 nM, or 100 nM Mlh1-Pms1 in a buffer containing 50 mM potassium acetate pH 7.9, 20 mM Tris-acetate, 10 mM magnesium acetate, and 100 μg/mL BSA for 10 minutes at room temperature. 2 units of T7 exonuclease was then added and reactions were incubated for 30 minutes at 37°C and stopped using 1% SDS, 14 mM EDTA, 0.96 units proteinase K (final concentrations) for 10 minutes. Reactions were then analyzed by 1% (w/v) agarose gel run at 100 V for 60 min, stained, visualized, and quantified as described above.

### Data Quantification and Statistical Analyses

All gels and TLC plates were imaged using a Sapphire Biomolecular Imager (Azure) and quantified using ImageJ software. In endonuclease experiments assayed by native agarose gel, nicking activity is quantified as the amount of DNA converted to nicked plasmid from supercoiled substrate. The amount of nicked or relaxed plasmid in negative controls was subtracted as background. For endonuclease experiments assayed by denaturing agarose gel, because Mlh1-Pms1 endonuclease activity is non-specific across substrate molecules, nicking activity is quantified as the amount of substrate DNA lost in reactions relative to negative controls. In electrophoretic mobility shift assays on plasmid DNA, because Mlh1-Pms1 binds to large DNA molecules as oligomeric complexes (10–12, 19, 35–37) and contorts their shape (12, 37), the amount of DNA bound was quantified as the loss of DNA in the substrate band relative to negative controls.

All experiments were performed in at least duplicate, with the number of replicates given for each experiment. When data is presented as a plot, the mean is shown with error bars representing the standard deviation between experiments.

## RESULTS

### ATP has opposing effects on Mlh1-Pms1’s affinity for DNA with and without breaks

Previous studies using *E. coli* MutL have shown that the protein recognizes 3’-resected DNA ends through an ATP-dependent mechanism (38). In *E. coli*, MutH initiates mismatch removal by nicking the newly replicated strand at hemi-methylated GATC sites, located on either the 5’ or the 3’ side of the mismatch. Since polymerases operate in the 3’ to 5’ direction, starting replication at a nick 3’ to the mismatch would not correct the mismatch but rather lead to DNA replication in the opposite direction from the mismatch. Thus, it was hypothesized that MutL’s ability to recognize 3’-resected ends could prevent improper nick processing by polymerases and promote mismatch removal by exonucleases or the UvrD helicase, which can remove the mismatch in either direction (38).

Eukaryotic mismatch repair does not rely on hemi-methylated GATC sites as strand discrimination markers, nor does it involve MutH as the endonuclease. Instead, eukaryotes employ a MutL homolog complex, Mlh1-Pms1 in yeast, as the endonuclease. *In vitro,* Mlh1-Pms1 displays PCNA-dependent strand discrimination and nicks plasmid substrates on the strand with a pre-existing discontinuity (14, 15). Given that these pre-existing discontinuities are necessary for strand bias *in vitro* and their maintenance is critical for efficient mismatch repair *in vivo* (22), we aimed to investigate whether eukaryotic Mlh1-Pms1 possesses the ability to recognize DNA discontinuities. To do this, we measured the relative affinities of Mlh1-Pms1 for oligonucleotide substrates that are intact or contain a single strand break constructed from radiolabeled oligonucleotides. Using electrophoretic mobility shift assays using increasing concentrations of yeast Mlh1-Pms1 at near physiological ionic strength (120 mM NaCl) and under conditions where we do not observe endonuclease activity (no PCNA is present), we observed near equivalent affinity for DNA with a discontinuity (K_d_ = 79 ± 3 nM) compared to intact 75-mer duplex (K_d_ = 96 ± 20 nM) (Figure S2). As described above, *E. coli* MutL displayed preferential binding for a 3’-resected DNA end in the presence of ATP (38). When we included ATP, there was no significant change in affinity compared to experiments without ATP for both the intact substrate (K_d_ = 79 ± 8 nM) and the discontinuous duplex (K_d_ = 77 ± 6 nM) (Figure S2). These data suggest that on small oligonucleotide substrates that support approximately two Mlh1-Pms1 copies per substrate, Mlh1-Pms1 does not specifically recognized DNA strand discontinuities.

Previous work has suggested that MLH endonucleases bind DNA cooperatively, where multiple proteins are present on a single DNA duplex, and that the protein functions as an oligomeric complex. These studies also suggest that formation of the oligomeric complex distorts the DNA substrate and interactions between oligomeric units in this assembly are necessary for activating the endonuclease activity (10–12, 19, 35, 36). Consistent with models for MLH oligomers, are findings that only large DNA substrates of at least ∼1 kb in size support efficient MLH endonuclease activity (11, 12) and DNA binding activities experience a sharp rise in efficiency when substrates are at least 500 bp in length (10). Because large DNA molecules are necessary to support MLH enzymatic activities and are the substate of choice for reconstituted mismatch repair reactions, plasmid substrates are a more relevant choice for Mlh1-Pms1 characterization studies than oligonucleotide substrates. Due to this, we measured Mlh1-Pms1’s affinity for large plasmid-based DNA molecules with and without discontinuities (Table 1, Figure 1, Figure S1, S3-4). We generated a range of circular plasmid substrates differing in the quantity of discontinuities, varying from zero to five single-strand breaks. We then measured the affinity of Mlh1-Pms1 for these substrates. Our findings revealed a decrease in the K_d_ value as the number of DNA breaks increase (Figure 1A-B, Figure S4), thereby affirming the binding selectivity of Mlh1-Pms1 for single-strand DNA breaks on substrates of a size that supports efficient DNA binding and endonuclease activity.

**Figure 1.**
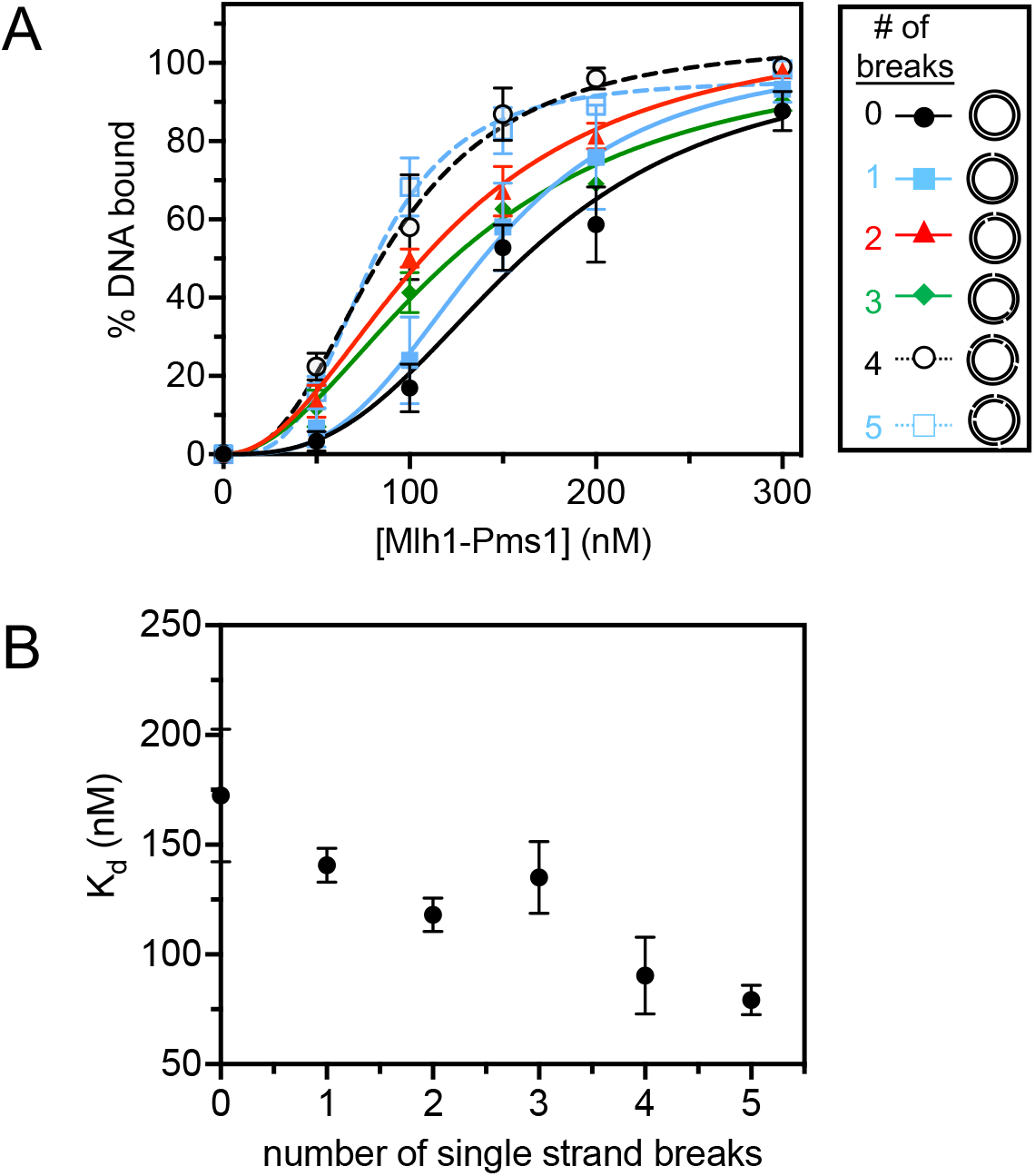
Mlh1-Pms1’s has higher affinity for DNA with single strand breaks than DNA without breaks. (A) Mlh1-Pms1 DNA binding curves on 2.7 kb DNA substrates that are fully relaxed or containing a variable number of single strand breaks. Single strand breaks were generated according to the Materials and Methods section. Mlh1-Pms1 was titrated to final concentrations: 0, 50, 100, 150, 200, or 300 nM. For substrates with 0, 2, 3, or 5 nicks, n = 3. For substrates with 1 or 4 nicks, n = 4. Data were fit to a sigmoidal function modeling cooperative binding. Hill coefficients were ∼3.0 for all curves. Representative primary data are in Figure S4. (B) K_d_ values were obtained from the binding data in A and plotted relative to the number of nicks.

**Table 1.** Mlh1-Pms1 has modestly higher affinity for DNA with single strand breaks in the presence of ATP. Mlh1-Pms1’s affinity for linearized plasmid either without discontinuities (intact DNA) or with four single strand (ss) breaks was measured by electrophoretic mobility shift assay. Single strand breaks were generated using a restriction nicking endonuclease from NEB. Representative primary data is in Figure S3. Where +, the concentration of ATP is 0.5 mM and the concentration of PCNA is 500 nM. K_d_ values were averaged and standard deviation is indicated. K_d_ values were averaged from n = 3 for all conditions except for intact DNA (-ATP, -PCNA) where n = 4. Standard deviations between experiments are indicated.

| DNA substrate | $K_d (\pm SD)$<br>(nM) | ATP | PCNA |
| --- | --- | --- | --- |
| Intact DNA | $140 \pm 17$ | - | - |
| | $210 \pm 13$ | + | - |
| | $210 \pm 10$ | - | + |
| | $250 \pm 28$ | + | + |
| DNA with ss breaks | $120 \pm 3$ | - | - |
| | $100 \pm 2$ | + | - |
| | $140 \pm 4$ | - | + |
| | $140 \pm 30$ | + | + |

In addition to Mlh1-Pms1 needing to form an oligomeric complex on DNA to be endonuclease active, PCNA is also required to activate the endonuclease activity. PCNA and DNA are also known to independently and synergistically enhance the ATPase activity of Mlh1-Pms1 (16, 28, 29). Due to this, we aimed to determine the effect of ATP and PCNA on Mlh1-Pms1’s affinity for single-strand breaks on large DNA molecules. To facilitate this, we used linearized plasmid as the DNA substrate which both supports *in vitro* endonuclease activity (11, 12) and allows for PCNA to load onto the free DNA ends without RFC. In these experiments, the sliding clamp nature of PCNA prevents measuring a stable PCNA-DNA complex. Thus, we are only able to measure the affinity of Mlh1-Pms1 for DNA. In DNA binding assays performed in the absence of PCNA, we found that on intact linear DNA that does not contain discontinuities, Mlh1-Pms1 binds with moderate affinity (K_d_ = 140 ± 17 nM). When ATP is added, the binding affinity is slightly reduced (K_d_ = 210 ± 13 nM), as has been previously observed (28, 29). Using linearized plasmid DNA that contains discontinuities, Mlh1-Pms1’s affinity (K_d_ = 120 ± 3 nM) is similar to what we measured on intact DNA, but when we included ATP, the affinity increased somewhat (K_d_ = 100 ± 2 nM), opposite to the effect observed on intact DNA (Table 1, Figure S3). These data suggest that when Mlh1-Pms1 binds to long DNA substrates that support oligomeric complexes, eukaryotic Mlh1-Pms1 selectively binds to DNA discontinuities in an ATP-dependent manner.

In the presence of PCNA, on intact DNA, we found that PCNA somewhat lowered the affinity of Mlh1-Pms1 for DNA (Table 1, compare K_d_ values of 140 ± 17 nM and 210 ± 10 nM, Figure S3) and the inclusion of both PCNA and ATP lowered the affinity of Mlh1-Pms1 for DNA slightly more, increasing the K_d_ value to 250 ± 28 nM. On DNA with single strand breaks, however, PCNA increased the K_d_ by a smaller margin and the combination of PCNA and ATP had little effect on Mlh1-Pms1’s affinity (Table 1, Figure S3). It should be noted that we performed these experiments on plasmid-based substrates to measure the behaviors of an Mlh1-Pms1 oligomer and its interactions with PCNA. However, because the substrate contains single-strand breaks, most of the DNA remains a continuous duplex, making the observed effects subtle. Although, only small changes in affinities between Mlh1-Pms1 and DNA were observed among the tested conditions, our data suggest that in the presence of DNA discontinuities, Mlh1-Pms1 does not interact proficiently with PCNA or use ATP hydrolysis to dissociate.

### Mlh1-Pms1’s ATPase activity loses PCNA-stimulation in the presence of DNA with discontinuities

Our observation showed that the affinity of Mlh1-Pms1 for DNA with single strand breaks remained largely unchanged in the presence of PCNA and ATP. Previous research has indicated that interactions between Mlh1-Pms1 and PCNA moderately stimulate Mlh1-Pms1’s ATPase (16, 28, 29). We therefore hypothesized that in the presence of DNA with single strand breaks, Mlh1-Pms1 should not effectively interact with PCNA, and we should observe minimal stimulation of Mlh1-Pms1’s ATPase activity. To test this, we performed an ATPase assay measuring the amount of ATP that is hydrolyzed by Mlh1-Pms1 with intact DNA or DNA that contains discontinuities. Under each condition we also included PCNA to see if ATP hydrolysis was stimulated (Figure 2A). In these experiments, and consistent with previous results, we found that in the presence of intact DNA, Mlh1-Pms1’s ATPase activity was stimulated by PCNA suggesting that Mlh1-Pms1’s ATPase is activated through PCNA interactions (Figure 2A first two columns, (16, 28, 29)). We then performed the experiment with DNA containing single strand breaks. In the presence of this substrate, we no longer observe a stimulatory effect of PCNA (Figure 2A last two columns). Similar to our DNA binding assay in Table 1, this suggests that Mlh1-Pms1 does not interact productively with PCNA on DNA containing single strand breaks, indicating that the DNA-break bound conformation of Mlh1-Pms1 is distinct from its intact DNA bound conformation.

**Figure 2.**
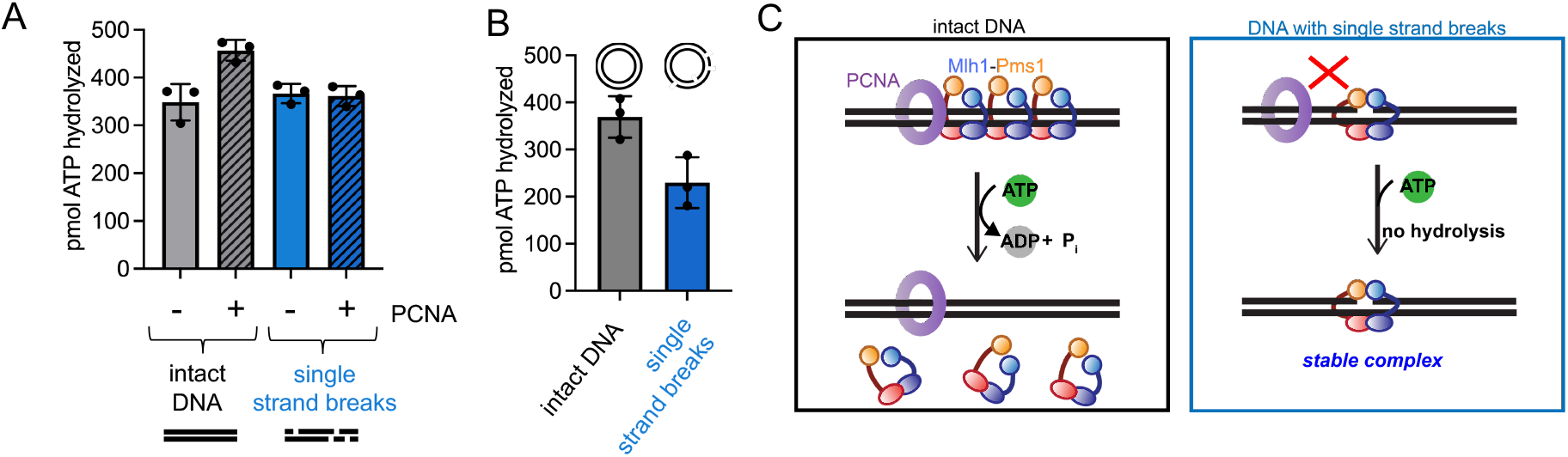
Mlh1-Pms1’s intrinsic ATPase activity loses PCNA stimulation in the presence of DNA breaks. (A) TLC ATPase assay measuring the amount of ATP hydrolyzed by Mlh1-Pms1 on linear 4.3 kb DNA. Grey bars represent intact linear 4.3 kb DNA (n = 3) and blue bars represent linearized 4.3 kb DNA with 4 single strand breaks (n = 3). Substrates were generated as described in Materials and Methods. Grey and blue bars with diagonal lines through them represent experiments containing PCNA (n = 3). (B) Grey bar represents percentage of ATP hydrolyzed on 4.3 kb relaxed, circular DNA without nicks (n = 3) and the blue bar represents circular 4.3 kb DNA containing 4 nicks (n = 3). 4.3 kb pBR322 was relaxed using topoisomerase I (NEB). Single strand breaks were made using Nt.BstNBI. (C) Model for the ATPase activity of Mlh1-Pms1 on intact DNA versus DNA containing single strand breaks.

In the experiment in Figure 2A we did not observe that Mlh1-Pms1 had a difference in ATPase efficiency in the presence of intact DNA relative to DNA with single strand breaks. In this assay, linearized plasmid was used as a DNA substrate to facilitate PCNA self-loading onto the free ends, so that we could perform the experiment in the absence of PCNA’s loader RFC, which is also an ATPase. Because the DNA is in a linear form, the termini of the DNA molecule may serve as additional specific binding sites for Mlh1-Pms1 preventing the observation of a difference in ATPase activity on these DNA substrates in the absence of PCNA. To directly test whether single strand breaks themselves affect Mlh1-Pms1’s ATPase efficiency, we measured the activity in the presence of circular DNA that is either intact or contains single strand breaks in the absence of PCNA (Figure 2B). Using these substrates, we observed that Mlh1-Pms1’s ATPase activity was slightly inhibited in the presence of single strand DNA breaks relative to intact DNA (Figure 2B). Because we also observed that the presence of ATP slightly enhanced Mlh1-Pms1’s affinity for DNA in the presence of breaks (Table 1, Figure S2), this suggests that Mlh1-Pms1 is in an ATP-bound state when recognizing these discontinuities. Since the protein’s ATPase is also no longer stimulated by PCNA on linear DNA with breaks, these data suggest that in the presence of DNA breaks, Mlh1-Pms1 likely binds to ATP to stabilize the complex but does not interact with PCNA or hydrolyze the bound ATP molecule (Figure 2C).

### Mlh1-Pms1’s endonuclease activity is inhibited on DNA with single strand breaks

If Mlh1-Pms1 does not interact with PCNA in the presence of single strand DNA breaks, then Mlh1-Pms1’s endonuclease activity, which requires interactions with PCNA, should be inhibited on DNA with single strand breaks compared to intact DNA. To test this hypothesis, we performed endonuclease assays on linear plasmid-based substrates at low ionic strength, which have been shown to support non-specific, but PCNA-dependent Mlh1-Pms1 endonuclease activity without a mismatch or Msh2-Msh6 (Figure 3). Furthermore, these substrates support self-loading of PCNA, so the reaction can be performed in the absence of RFC to prevent preferential loading at sites of single strand breaks. We monitored endonuclease activity on these substrates using denaturing agarose gels, measuring the degradation of DNA due to Mlh1-Pms1’s non-specific nicking activity. On intact DNA, we observed robust endonuclease activity that increased as a function of Mlh1-Pms1 concentration (Figure 3A). When we performed this experiment with DNA with one single strand break, we found that Mlh1-Pms1’s endonuclease activity was slightly inhibited compared to the intact substrate (Figure 3B, D). When we increased the number of single strand breaks to four, we found that Mlh1-Pms1’s endonuclease was even further inhibited (Figure 3C, D). This further suggests that in the presence of DNA breaks, Mlh1-Pms1 can recognize the DNA break but is unable to interact with PCNA in a conformation that promotes ATPase or endonuclease enzymatic activities (Figure 3E).

**Figure 3.**
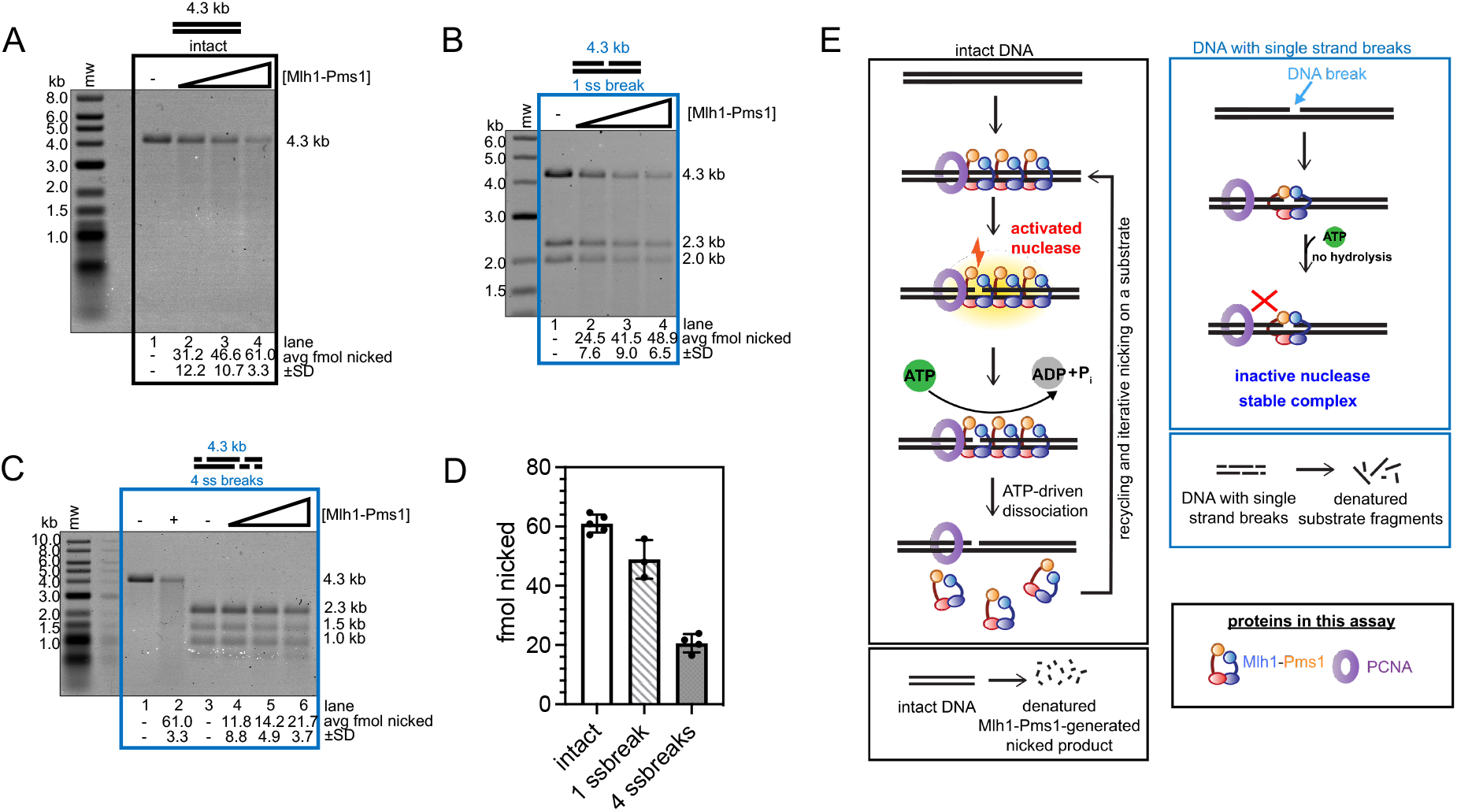
Mlh1-Pms1’s endonuclease activity on DNA is inhibited in the presence of single strand breaks. Endonuclease assays performed as described in the Materials and Methods section and analyzed by denaturing gel electrophoresis. All reactions contain 0.5 mM ATP and 500 nM PCNA. Because endonuclease activity is non-specific on substrates in this figure, the amount of DNA nicked is measured as the degradation of DNA bands relative to negative controls. Where indicated, 1 kb plus ladder (mw) (NEB) was used to measure starting material fragment sizes. Sizes indicated to the right of the gel are lengths of initial DNA fragments that form the substrates. (A) Intact linear 4.3 kb (n=4) substrates incubated with increasing concentrations of Mlh1-Pms1. Lanes 1-4 contain 0 (n=5), 50, 100, and 200 nM (n=5) Mlh1-Pms1, respectively. (B) Linearized 4.3 kb (n = 3) DNA substrates containing with one single strand break generated with Nt.BspQI were incubated with increasing amounts of Mlh1-Pms1. Lanes 1-4 contain the same concentrations of Mlh1-Pms1 as in A. (C) Linearized 4.3 kb substrates (n = 5) containing four single strand breaks generated with Nt.BstNBI were incubated with increasing amounts of Mlh1-Pms1. Lanes 1 and 2 are negative and positive controls (where +, Mlh1-Pms1 is 200 nM) with continuous 4.3 kb substrates, respectively. Lanes 3-6 contain the same concentrations and A and B. (D) Comparison of the amount of DNA nicked, in fmol, using 200 nM Mlh1-Pms1 among the 4.3 kb substrates. (E) Model for Mlh1-Pms1 activities in this figure.

Similarly, oligomers of the meiotic homolog Mlh1-Mlh3 have previously exhibited similar activities to what we observe for Mlh1-Pms1. Mlh1-Mlh3 has been shown to have moderately high affinity for substrates with insertion loops (11). Although that protein had increased affinity for those substrates, the DNA loop was observed to disrupt the oligomeric assembly and partially inhibit endonuclease activity. Because Mlh1-Mlh3 and Mlh1-Pms1 are homologs and share a subunit, we hypothesize that single strand breaks may play a similar role and disrupt or terminate an Mlh1-Pms1 oligomer. As discussed previously, Mlh1-Pms1 binds to substrates ≥500 bp and requires substrates to be larger than 1 kb for efficient endonuclease activity. Although, the duplex nature of DNA maintains the molecular architecture of the model plasmid substrate, single strand breaks may disrupt or terminate an Mlh1-Pms1 oligomer, making the substrate behave more like consecutive small DNA fragments of sizes that do not support oligomeric complexes and endonuclease activity (Figure S1).

### Mlh1-Pms1 uses the ATPase activity in Mlh1 to iteratively nick DNA

In the assays in Figure 3, endonuclease activity presents as the apparent degradation of DNA substrate due to the non-specific nature of the endonuclease activity and likely repeated nicking events on the same substrate. Although never directly established, the repetitive nicking by Mlh1-Pms1 is hypothesized to result from recycling of Mlh1-Pms1 on DNA promoted through the protein’s ATPase (6, 8, 9, 28). This is supported by the observed lower affinity between Mlh1-Pms1 and DNA in the presence of ATP presented here (Table 1, Figure S2, Figure S3). Iterative rounds of endonuclease activity induced by Mlh1-Pms1 ATPase activity is also supported by the stimulation of Mlh1-Pms1’s ATPase by DNA, the observed large conformational shifts in the protein in the presence of ATP, and single molecule and biochemical work observing fewer nicking events with mutants that have deletions in the intrinsically disordered regions that also show ATPase defects (16, 28–30, 39, 40).

To directly test for Mlh1-Pms1’s ability to use its ATPase function to recycle and iteratively nick the same DNA substrate, we generated a mutant version of Mlh1-Pms1 that is defective in ATP binding and hydrolysis in both subunits, mlh1N35A-pms1N34A (25, 27). In *in vivo* experiments measuring mismatch repair efficiency in bakers’ yeast, both a *mlh1N35A* and a *pms1N34A* strain displayed a mutation rates similar to null strains (27). In biochemical characterization assays for these mutants, the amino-terminal ATPase domains of either Mlh1 or Pms1 were isolated as truncations of either wild-type, mlh1N35A, or pms1N34A mutants. In these experiments, both the isolated amino terminal domain of Mlh1 with the N35A mutation and the isolated amino terminal domain of Pms1 with the N34A mutation were deficient in both ATP binding and hydrolysis when measured directly (27).

Using the mlh1N35A-pms1N34A ATPase mutant, we performed endonuclease assays to measure the total number of DNA molecules that are nicked by the endonuclease. In this experiment, non-specific endonuclease activity is assessed as conversion of supercoiled DNA to nicked circular product. In this assay, we observed that the mlh1N35A-pms1N34A ATPase mutant exhibited a mild defect in endonuclease activity (Figure 4A, compare lanes 2-5 with 7-10). As stated, this defect in activity is measured as conversion of supercoiled to nicked DNA, but whether a substrate was nicked multiple times cannot be determined.

**Figure 4.**
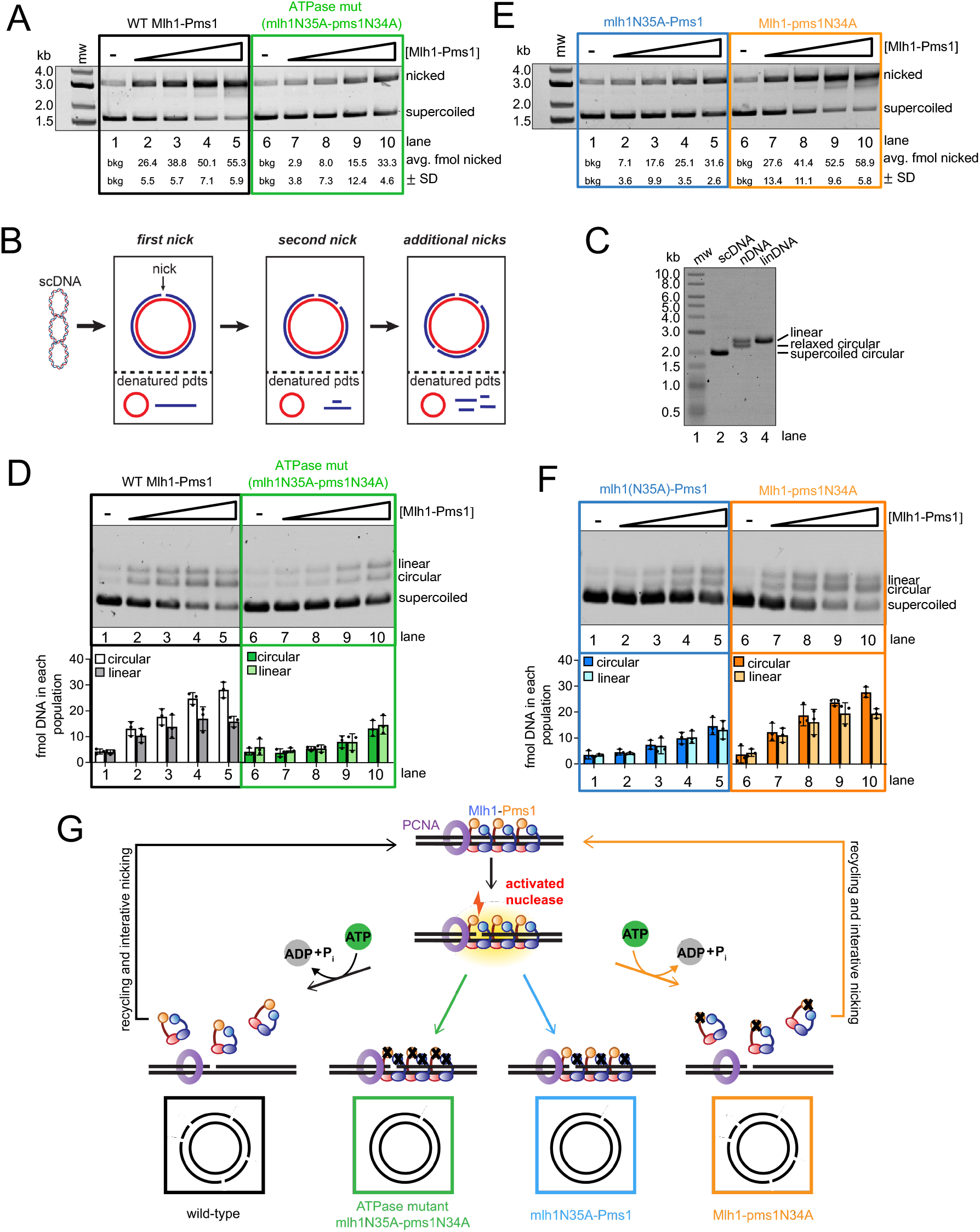
The ATPase activity in the Mlh1 subunit drives Mlh1-Pms1 endonuclease recycling on DNA. (A) Endonuclease assay measuring population of 2.7 kb supercoiled DNA (135 ng) substrates that are nicked at least once by either wild-type Mlh1-Pms1 or mlh1(N35A)-pms1(N34A) ATPase mutant. Either 0, 25, 50, 100, 200 nM of wild-type Mlh1-Pms1 or ATPase mutant was incubated with the supercoiled DNA, along with 500 nM PCNA, and 100 nM RFC (n = 3). (B) Schematic for denaturing analysis of the material in panel A. In denaturing gels, iterative nicking is detectable as a loss of signal and nicking is expected to be biased towards the strand that is initially nicked due to the nick being used as a preferential site for additional PCNA loading which can impart strand bias (see Results section). (C) Control denaturing gel demonstrating migration of possible DNA products produced in this assay. Lane 2 contains supercoiled 2.7 kb DNA (pUC18), lane 3 contains 2.7 kb DNA which contains one nick (Nt.BspQI), lane 4 contains 2.7 kb DNA linearized with HindIII. (D) Denaturing analysis of material in A (n = 3). (E) The number of supercoiled plasmids nicked by wither mlh1N35A-Pms1 or Mlh1-pms1N34A (n = 3) was measured identically to how performed in A. (F) Denaturing analysis of material in E (n = 3). (G) Model for Mlh1-Pms1 activities in this figure.

To determine if the mlh1N35A-pms1N34A ATPase mutant is defective in iteratively nicking the same DNA substrate, we analyzed the reaction products from the assay in Figure 4A in a denaturing gel system also used in Figure 3. In a denaturing gel, if a supercoiled substrate is nicked once, the expected reaction products are a circular DNA strand and a linear single strand. If the same strand is nicked multiple times, the expected products are a circular DNA strand and smaller fragments of the linear single strand that was originally nicked (Figure 4B). This preferential degradation of the DNA that contains the first nick is due to strand discrimination, imparted by PCNA which is preferentially loaded onto nicked DNA by RFC in a specific orientation (41). Using this system to assay for repetitive nicking, we determined that although the wild-type Mlh1-Pms1 was able to iteratively nick each plasmid substrate, with repetitive nicking biased towards the same strand (Figure 4C for migration marker, Figure 4D, lanes 1-5) (8, 9, 28), the mlh1N35A-pms1N34A ATPase mutant was defective in this activity (Figure 4D, lanes 6-10). Although suggested by data in Figure 1 and Table 1, these data directly suggest that Mlh1-Pms1 uses its ATPase activity to recognize breaks in DNA, likely including the nicks that it generates, promoting ATP hydrolysis and recycling to promote iterative nicking.

Previous biochemical research indicates that MutL homolog subunits exhibit asymmetric behavior when dimerized, whether forming a MutL homodimer with bacterial subunits or eukaryotic heterodimers (25, 27, 29). Specifically, for yeast Mlh1-Pms1, studies have shown that the truncated Mlh1 ATPase domain has a lower K_m_ compared to the truncated Pms1 ATPase domain. Additionally, in kinetic ATPase assays, the Pms1 ATPase domain exhibits a higher k_cat_ value than the Mlh1 ATPase domain (27).

Given our observed mechanistic link between total ATPase function of Mlh1-Pms1 and the ability of the protein to interact with PCNA and recycle on DNA, we aimed to identify which subunit’s ATPase activity primarily drives recycling, particularly considering Pms1’s presumed role as the primary PCNA-interacting subunit. To address this, we expressed and purified mutant variants: one with impaired ATPase activity solely in the Mlh1 subunit (mlh1N35A-Pms1), and another with deficiency solely in the Pms1 subunit (Mlh1-pms1N34A). We then measured endonuclease activity in a population-based assay to determine the total number of DNA molecules nicked by the Mlh1-Pms1 ATPase variants. Our findings revealed that the Mlh1-pms1N34A mutant displayed activity comparable to the wild type. In contrast, the mlh1N35A-Pms1 mutant exhibited slightly inhibited activity, with nicking efficiencies similar to those of the mlh1N35A-pms1N34A mutant (see Figure 4E compared to Figure 4A). By analyzing the reaction products using the denaturing analysis in Figure 4B-C, which allows us to observe iterative nicking events, we similarly found that the mlh1N35A-Pms1 mutant was inhibited for promoting multiple rounds of endonuclease activity directed to the same substrate molecule. The Mlh1-pms1N34A mutant was proficient in this activity (Figure 4F). These observations indicate that the Mlh1 subunit’s ATPase activity primarily facilitates Mlh1-Pms1 recycling after DNA nicking. Additionally, the data suggest that the Pms1 subunit, housing the endonuclease motif, likely recognizes its own endonuclease product and stabilizes the complex. This hypothesis is supported by the fact that mlh1N35A-Pms1 complexes do not appear to dissociate after nicking DNA, suggesting that Pms1 likely plays a significant role in recognizing the endonuclease product (Figure 4G). These data are also in agreement with literature where Mlh1 has higher affinity for DNA compared to Pms1, suggesting that Mlh1 controls the recycling of the protein after endonuclease activity to iteratively nick DNA (27).

### Mutants defective in ATPase activity are trapped after nicking DNA

The motifs facilitating ATPase activity in MutL homologs are highly conserved across organisms (26). Consistent with an important biological significance, missense mutations in the ATPase region of human MLH1 have been identified in patients with Lynch syndrome, a hereditary cancer syndrome associated with mismatch repair defects (42–44). Included in this set are mutations to residue N38 in human MLH1 which aligns with N35 in yeast Mlh1 (Figure S6A). Mutations adjacent to the ATPase motif also have been shown to affect the cellular localization of mammalian MLH1 and are associated with breast cancers (45). Because the ATPase site in Mlh1 is biologically significant across organisms (27), and our findings in Figure 4 indicate that an Mlh1-Pms1 ATPase mutant retains functional endonuclease activity, despite failures to recycle, we aimed to elucidate the consequences of impaired recycling to better explain *in vivo* defects.

To determine the fate of the mlh1N35A-pms1N34A mutant on DNA, we conducted a non-specific nicking reaction on supercoiled DNA, allowing either the wild-type or the ATPase mutant protein to nick the DNA. Subsequently, T7 exonuclease was introduced to degrade DNA in the 5’ to 3’ direction, initiating at nicks generated by Mlh1-Pms1. We found that with the wild-type protein, T7 exonuclease nearly completely degraded the DNA molecules nicked by wild-type Mlh1-Pms1 (Figure 5A and C). This suggests that after nicking the DNA, Mlh1-Pms1 could recycle and dissociate from the nicks (Figure 5D). In contrast, reactions with the ATPase mutant showed inhibited degradation by the exonuclease (Figure 5B-C). These results indicate that the mlh1N35A-pms1N34A ATPase mutant not only fails to recycle *in vitro* but also becomes trapped at DNA nicks, protecting the entire substrate from T7 degradation (Figure 5D). Despite its competence in endonuclease activity, the protein remains tethered at the nick site, providing a possible explanation for the observed defects both in yeast genetic assays and the association with Lynch syndrome (25, 27, 43, 44).

**Figure 5.**
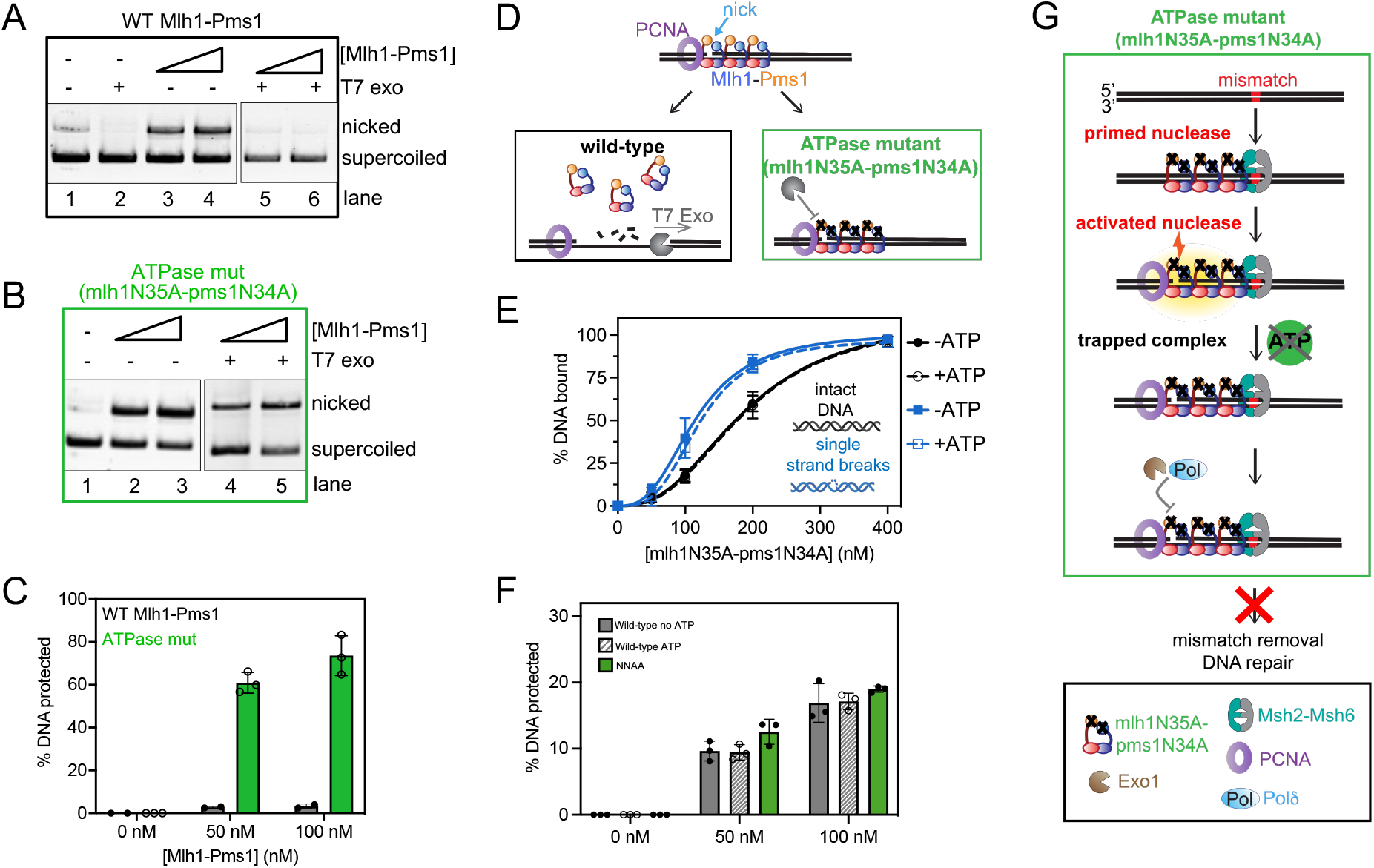
An Mlh1-Pms1 ATPase mutant is trapped at DNA nicks. (A-B) Either 0, 50, or 100 nM of wild-type Mlh1-Pms1 or mlh1N35A-pms1N34A ATPase mutant was incubated with supercoiled DNA where indicated in the presence of 500 nM of PCNA, 100 nM RFC, and 0.25 mM ATP under conditions that promote endonuclease activity. After incubation, T7 exonuclease was added where + indicated that degrades the substrate in the absence of Mlh1-Pms1 (Figure S5). Reaction products were analyzed by agarose gel. (C) The amount of DNA protected from T7 exonuclease degradation was calculated by quantifying the amount of nicked DNA for each Mlh1-Pms1 concentration in the absence of T7 exonuclease relative to the amount that remains after treatment with T7 exonuclease. For wild-type Mlh1-Pms1 n = 2, for the mlh1N35A-pms1N34A ATPase mutant n = 3. (D) Model for Mlh1-Pms1 activities in this assay. (E) The mlh1N35A-pms1N34A ATPase mutant retains its affinity for single strand breaks but is affinity is unaffected by ATP. Relative DNA affinities were measured by electrophoretic mobility shift assay as described in the Materials and Methods. (F) The amount of DNA protected from T7 exonuclease degradation was measured as in A-C using circular DNA with a single pre-existing nick generated by a restriction nicking endonuclease. (G) Model for mlh1N35A-pms1N34A ATPase mutant’s defects in mismatch repair *in vivo*.

To determine if the apparent trapping effect was simply due to mlh1N35A-pms1N34A having higher affinity for DNA breaks than the wild-type protein, we measured the relative affinities of the mlh1N35A-pms1N34A ATPase mutant for DNA with and without breaks (Figure 5E). We determined that the mlh1N35A-pms1N34A ATPase mutant bound to intact DNA with a K_d_ of 190 ± 20 nM and DNA with breaks with a K_d_ of 120 ± 10 nM, which are near wild-type affinities as measured in Table 1. When we included ATP in the assay, we observed near equivalent K_d_ values (190 ± 40 nM for intact DNA and 120 ± 6 nM for DNA with breaks) with experiments without ATP, consistent with the ATPase defect in the protein. This suggests that the failure to dissociate from nicked DNA product in Figure 5A-C is due to a failure in the ATPase activity to promote recycling. It also suggests distinct interactions between Mlh1-Pms1 and its own endonuclease product and pre-existing strand discontinuities.

We aimed to discern whether the trapping of the mlh1N35A-pms1N34A ATPase mutant at a DNA nick resulted from a specific conformation post-DNA incision, or if pre-existing single-strand breaks in the DNA induced a similar effect. We hypothesize that the wild-type Mlh1-Pms1 protein can access a conformation after nicking DNA that promotes ATP binding and hydrolysis stimulated through PCNA interactions that the mlh1N35A-pms1N34A ATPase mutant does not access. To test this, we generated a DNA substrate with a single pre-existing single strand break and pre-bound either the wild-type or mlh1N35A-pms1N34A ATPase mutant protein to the DNA. We omitted RFC and PCNA from the reaction to not induce endonuclease activity. We then added an amount of T7 exonuclease that in the absence of Mlh1-Pms1 degrades the entirety of the nicked strand (Figure S5). Using the DNA with the pre-existing break, we observed that both the wild-type and the mlh1N35A-pms1N34A ATPase mutant protein protected the DNA from T7 exonuclease degradation to similar extents (Figure 5F). It should be noted that pre-existing single-strand breaks in the DNA were introduced using commercially available restriction nicking endonucleases, which produce discontinuities containing a 3’-hydroxyl group and a 5’-phosphate group. Mlh1-Pms1 likely makes nicks with identical chemistries given that these nicks serve as substrates for polymerases and exonucleases in mismatch repair. Taking this inference into account along with the observation that the mlh1N35A-pms1N34A ATPase mutant and wild-type Mlh1-Pms1 protect pre-existing breaks to similar extents, the difference in protection observed in Figure 5A-C is likely caused by Mlh1-Pms1 accessing a specific conformation after nicking the DNA through PCNA activation and not simply through recognizing the DNA discontinuity. How Mlh1-Pms1 treats the nick it makes versus a pre-existing nick are therefore not equivalent and represent two modes of the protein (see Discussion). Mlh1-Pms1 uses ATP to both recognize and dissociate from the nick it makes when complexed with PCNA post-DNA incision. However, ATP binding is not required to recognize a pre-existing nick but can be used to stabilize the complex on DNA. Together these data suggest that defects observed for mlh1N35A-pms1N34A ATPase mutants *in vivo* (27) are caused by the protein making a nick to initiate mismatch removal, but failing to dissociate from the site preventing mismatch removal and allowing both the mismatch and the nick to persist (Figure 5G).

## DISCUSSION

We have observed that the eukaryotic Mlh1-Pms1 mismatch repair endonuclease oligomeric complex recognizes single strand breaks in DNA and that this affinity increases modestly in the presence of ATP (Table 1, Figure 1, Figure S2-S4). The fact that we could only see this effect on large substrates where Mlh1-Pms1 displays endonuclease activity and is expected to form an oligomeric complex suggests that either an oligomeric complex is needed to recognize a pre-existing nick or the Mlh1-Pms1 copy or copies that recognize this site are influenced by Mlh1-Pms1 oligomers elsewhere on the substrate. In addition to this, we also observed that in the presence of DNA breaks, Mlh1-Pms1 loses PCNA-dependent activities, including PCNA-stimulated ATPase and endonuclease activities (Figure 2-3). When we generated an Mlh1-Pms1 mutant that was defective in ATPase activity in both subunits, we determined that, although the protein could interact with PCNA and be activated to nick DNA substrates non-specifically, it was defective in recycling and iteratively nicking the DNA (Figure 4A-D). Upon generating ATPase mutants in Mlh1 and Pms1 subunits independently, we found that this effect was driven by the Mlh1 subunit (Figure 4E-F). Using an exonuclease protection assay, we determined that the Mlh1-Pms1 ATPase mutant remained at DNA nick sites whereas the wild-type protein dissociated (Figure 5). The trapped state of the Mlh1-Pms1 ATPase mutant suggests that the defect in iterative nicking is likely due to a failure to dissociate from the nicked DNA product and reset.

Our data suggest that Mlh1-Pms1 can access two distinct modes on nicked DNA which use ATP uniquely depending on whether Mlh1-Pms1 itself generated the nick or if the nick was pre-existing and needed for strand discrimination to facilitate proper mismatch repair. In post-replicative mismatch repair, Mlh1-Pms1 is recruited to an error-containing substrate that is intact in the vicinity of the mismatch. Mlh1-Pms1 is then activated to nick the nascent DNA strand through interactions with PCNA. Interactions with PCNA, then promote ATP binding and hydrolysis driven by the Mlh1 subunit, which our data suggest triggers DNA dissociation and allows proteins that remove the mismatch to have access to the DNA nick (Figure 6, Mode 1, left side).

**Figure 6.**
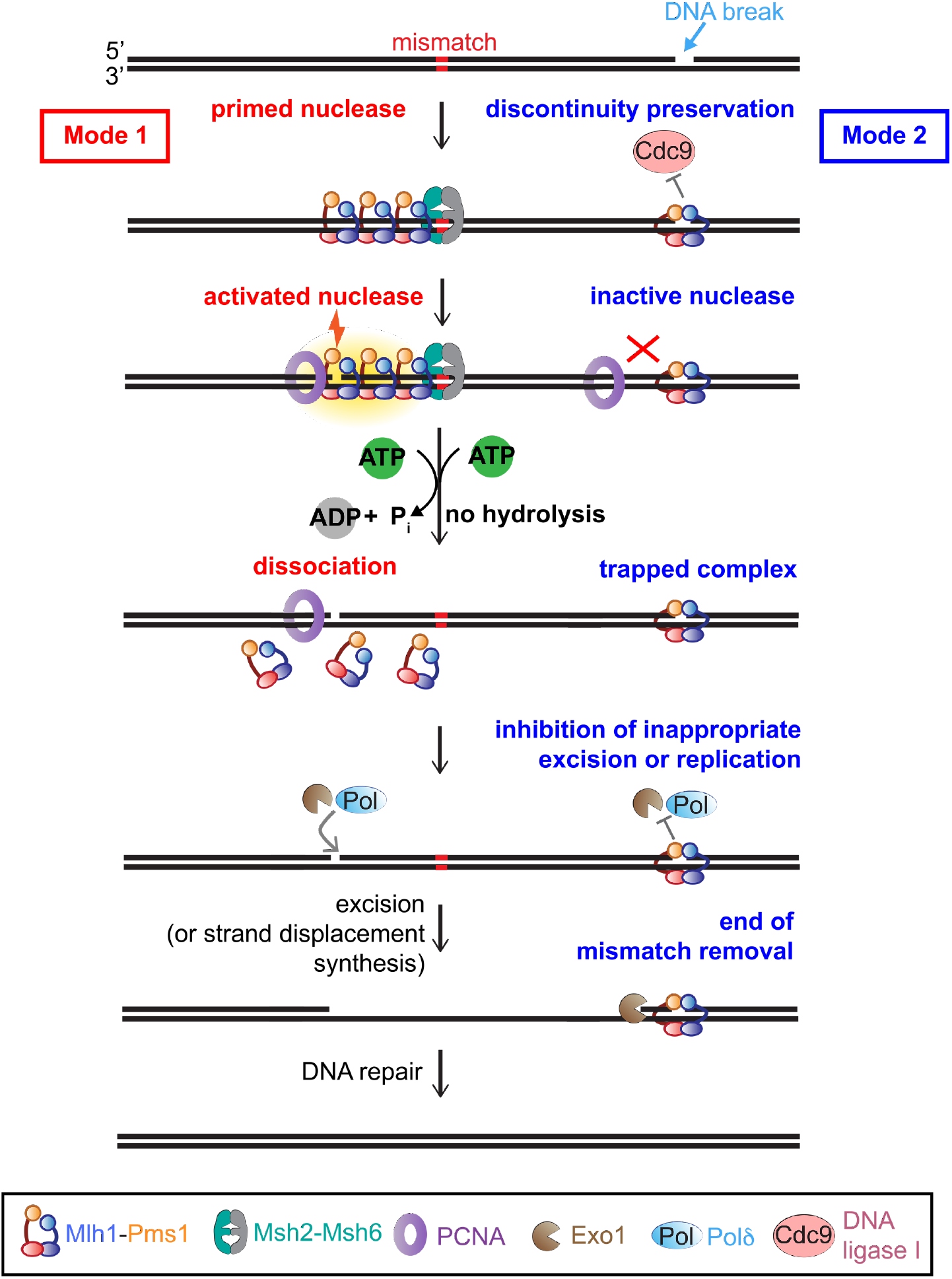
Model for the role of Mlh1-Pms1 single strand break recognition and ATPase activities *in vivo*. Mlh1-Pms1 uses the presence or absence of discontinuities in the DNA and its ATPase activities to toggle between activity modes. In the absence of discontinuities, Mlh1-Pms1 can prime its endonuclease activity (Mode 1, red) to interact with PCNA, nick DNA, hydrolyze ATP through PCNA interactions, and dissociate or recycle to signal mismatch removal. In the presence of DNA breaks or discontinuities, Mlh1-Pms1 can recognize the break (Mode 2, blue) and protect the break from ligation. The pre-existing nick-bound Mlh1-Pms1 does not efficiently interact with PCNA, which inhibits its endonuclease activity and attenuates its ATPase activity. This recognition can also protect breaks in the DNA that are not sites for mismatch removal, prevent to formation of more breaks introduced by Mlh1-Pms1 and may also serve as a termination point for mismatch removal. See Discussion.

As a second mode, our data suggest that pre-existing DNA discontinuities that are byproducts of replication and other repair processes (1, 20, 21) can recruit Mlh1-Pms1 in the absence of Msh2-Msh6 or a mismatch. The Mlh1-Pms1 protein recruited by these single strand breaks is prevented from introducing additional strand breaks and dissociating by its failure to productively interact with PCNA and hydrolyze ATP. Instead, ATP binding is used to stabilize the complex at the break, trapping it. We hypothesize that this stabilization serves as a protection mechanism preventing nick ligation by Cdc9 (DNA ligase I) and preventing Pol8 or Exo1 from inappropriately processing the break that is not adjacent to a mismatch, as suggested for *E. coli* MutL’s affinity for 3’-recessed ends (38). Mlh1-Pms1’s specificity for pre-existing breaks may also serve as a termination mechanism for mismatch removal initiating at the Mlh1-Pms1-generated nick (Figure 6, Mode 2, right side).

As mentioned earlier, in post-replicative mismatch repair, the nascent DNA strand can have discontinuities on the lagging strand between Okazaki fragments and on the leading strand from non-processive replication or other repair processes (1, 20, 21). Work in yeast has demonstrated that the maintenance of discontinuities on nascent DNA is necessary for efficient mismatch repair *in vivo* (22) and work in yeast, human, and *Xenopus* systems demonstrates their importance *in vitro* and suggests that they serve as strand bias landmarks by acting as PCNA loading sites (4, 9, 14, 15, 17, 18, 41). In these models PCNA ultimately dictates strand discrimination, but the mechanistic details of this communication channel are not established. Precisely how PCNA retains bias after being loaded at pre-existing nicks is unclear. Additionally, it is unclear *in vivo* whether the PCNA that activates Mlh1-Pms1 in mismatch repair is loaded distinctly for repair or deposited by the replication machinery since Mlh1-Pms1’s activities are temporally and spatially separated from DNA replication (19). Despite these questions for how strand discontinuities regulate mismatch repair, retaining these discontinuities as signals is important for mutation avoidance (22) and our data suggest a mechanism for how Mlh1-Pms1 protects these breaks from ligation or inappropriate processing during repair.

Our models suggest that ATP plays two distinct and critical roles in Mlh1-Pms1 function. Consistent with this, in *in vivo* assays in *S. cerevisiae* measuring mismatch repair, mutants that are defective to ATP hydrolysis have null-like phenotypes (27). Our data measuring the fate of an ATPase mutant on DNA after endonuclease activities suggests that these mutants remain trapped at DNA nick sites, preserving the DNA damage similar to how PARP proteins can become trapped at DNA damage sites in the presence of PARP inhibitor (46, 47). This could account for the null phenotype *in vivo* in bakers’ yeast. Interestingly, although both the *mlh1-N35A* and the *pms1-N34A* (reported as pms1-N65A due to numbering from a different start codon) strains gave phenotypes similar to *mlh1Δ* and *pms1Δ* haploid strains, in experiments in diploid heterozygous strains, although an *MLH1/mlh1Δ* strain has a mutation rate only slightly above wild-type, an *MLH1/mlh1-N35A* strain has a rate 480-times the wild-type, suggesting that the mlh1-N35A protein may be poisonous to the pathway (27). This is consistent with our data which suggest that defects in Mlh1-Pms1 ATPase activity create trapped complexes on DNA after endonuclease activity. Failures in Mlh1-Pms1 turnover are driven largely by failures in Mlh1’s ATPase activity which is required for Mlh1-Pms1 to dissociate from the nick it makes and recycle on the DNA for iterative rounds of nicking.

The ATPase motifs in Mlh1 and Pms1 are highly conserved across organisms (Figure S6A). Based on structural work in *E. coli,* the N35 residue in the Mlh1 ATPase site and the N34 residue in the Pms1 subunit’s ATPase site are used to coordinate a divalent metal that is necessary to stabilize the negative charges on ATP (26) (Figure S6B). By mutating this residue and eliminating ATP binding and subsequent hydrolysis, we demonstrated a link between the ATPase activity, product recognition, and recycling. This residue in particular is highly conserved among MutL homologs and missense mutations to this residue (N38 in human) among others adjacent to the ATPase site are associated with Lynch syndrome (Figure S6) (43, 44). Our data suggest that a mechanistic explanation for this disease state may be due to this protein making a nick adjacent to a mismatch but then failing to dissociate from DNA (Figure 5G). In this scenario, not only does the mismatch persist, but additional DNA damage has been introduced by human MLH1-PMS2’s endonuclease activity, and the persistence of an MLH1-PMS2 complex may cause additional replication stress and trigger additional instability.

## DATA AVAILABILITY

This study includes no data deposited in external repositories.

## SUPPLEMENTARY DATA

A separate file is available with supplementary data.

## ACKNOWLEDGEMENTS

We thank all members of the Manhart lab and Eric Alani for helpful discussions about the work in this manuscript. We also thank the Department of Chemistry and the College of Science and Technology at Temple University for their support. The content of this work is solely the responsibility of the authors and does not necessarily represent the official views of the National Institutes of Health. The funders had no role in the study design, data collection and analysis, decision to publish, or preparation of the manuscript.

## FUNDING

National Institute of General Medical Sciences of the National Institutes of Health [R35GM142651 to J.M.P., S.J.W., and C.M.M.]; College of Science and Technology at Temple University (to S.J.W. and Y.S.S., in part). Funding for open access charge: National Institutes of Health.

## CONFLICT OF INTEREST

None declared.

## Notes

### Competing Interest Statement

The authors have declared no competing interest.

