## Supplementary Material for "The Mlh1-Pms1 endonuclease uses ATP to preserve DNA discontinuities as strand discrimination signals to facilitate mismatch repair"

† Joint Authors

### SUPPLEMENTARY TABLE

#### Table S1

| Oligonucleotide Name | Sequence (5' to 3') | Purpose |
|---|---|---|
| CMO175 | AATGATGGAGGCGTCCATCGATGCG | Forward primer to generate N35A mutation in *MLH1* |
| CMO176 | TCTTTGAGAGCATTTACG | Reverse primer to generate N35A mutation in *MLH1* |
| CMO177 | ACTCGTTGATGCGAGTATAGATGCG | Forward primer to generate N34A mutation in *PMS1* |
| CMO178 | TCTTTCACTGCAGTTGTTAAG | Reverse primer to generate N34A mutation in *PMS1* |
| CMO168 | TGGGTCAACGTGAGCAAAGATGTCCTAGCAAGTCAGAATTC GGTAGCGTGACGTGACCTGCTAGACTTAGGTCGA | Complement to CMO169 to create continuous 75-mer duplex |
| CMO169 | TCGACCTAAGTCTAGCAGGTCACGTCACGCTACCGAATTCT GACTTGCTAGGACATCTTTGCTCACGTTGACCCA | 75-mer substrate, 5' radiolabeled |
| CMO340 | TGGGTCAACGTGAGCAAAGATGTCCTAGCAAGTCAGA | 37-mer complement to 3' end of CMO169 used in discontinuous oligonucleotide |
| CMO341 | ATTCGGTAGCGTGACGTGACCTGCTAGACTTAGGTCGA | 38-mer complement to CMO169 used in discontinuous substrate |

**Table S1. Oligonucleotides used in this study.** Primers CMO175-178 used to generate mutations to abrogate ATPase activity in Mlh1 and Pms1 subunits**.** Q5 mutagenesis (NEB) was performed using pMH1 and pMH8 as templates (see Materials and Methods) (31). CMO168-CMO170 and CMO184 were used to construct oligonucleotide substrates used to measure Mlh1-Pms1 DNA binding in Figure S2.

### SUPPLEMENTARY FIGURES

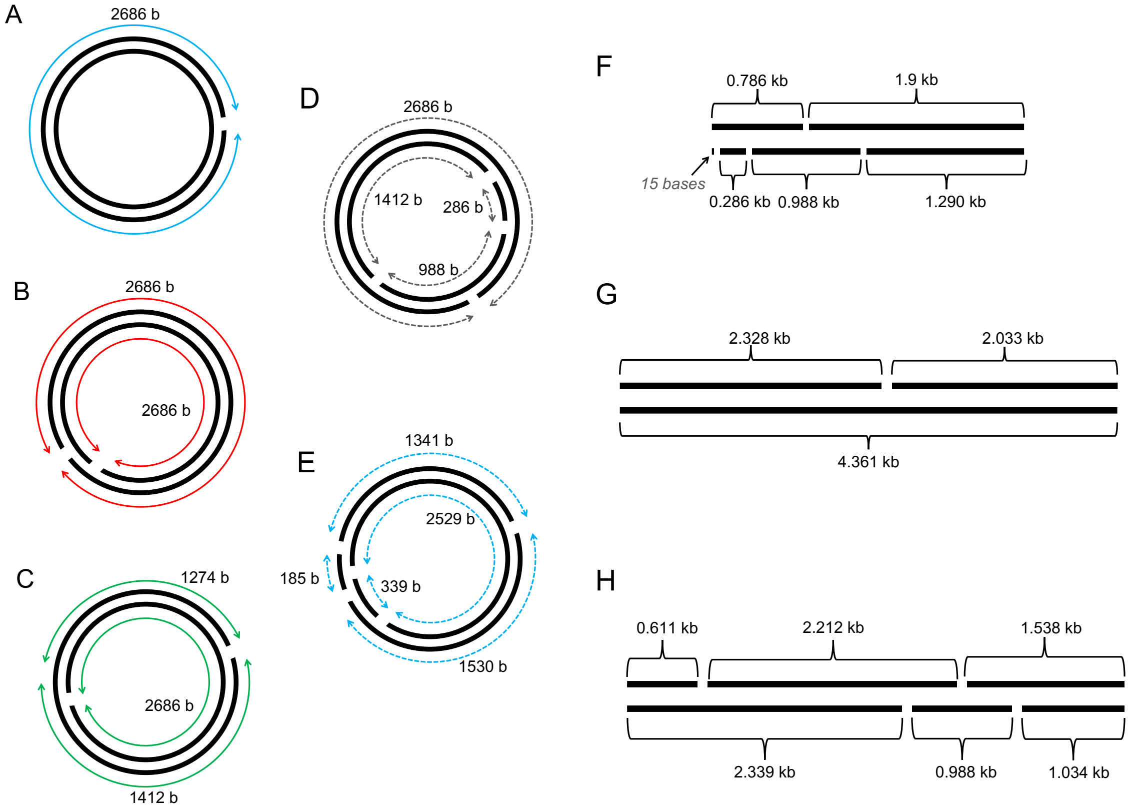


**Figure S1. Plasmid-based substrates used in this study**. (A-E) Location of sites for pre-existing nicks on circular plasmids (2.7 kb) used in experiments where circular substrates with breaks are used. Substrates contain between 1 and 5 pre-existing discontinuities introduced using restriction nicking endonucleases as described in the Materials and Methods. Distances and spacing between nicks are indicated on the maps in number of bases (b). (F-H) Location of sites for pre-existing nicks on linearized plasmids (2.7 kb in F, 4.3 kb in G-H) used in experiments where DNA was in linear form to facilitate PCNA self-loading. Distances and spacing between nicks are indicated on the maps in number of kilobases (kb).

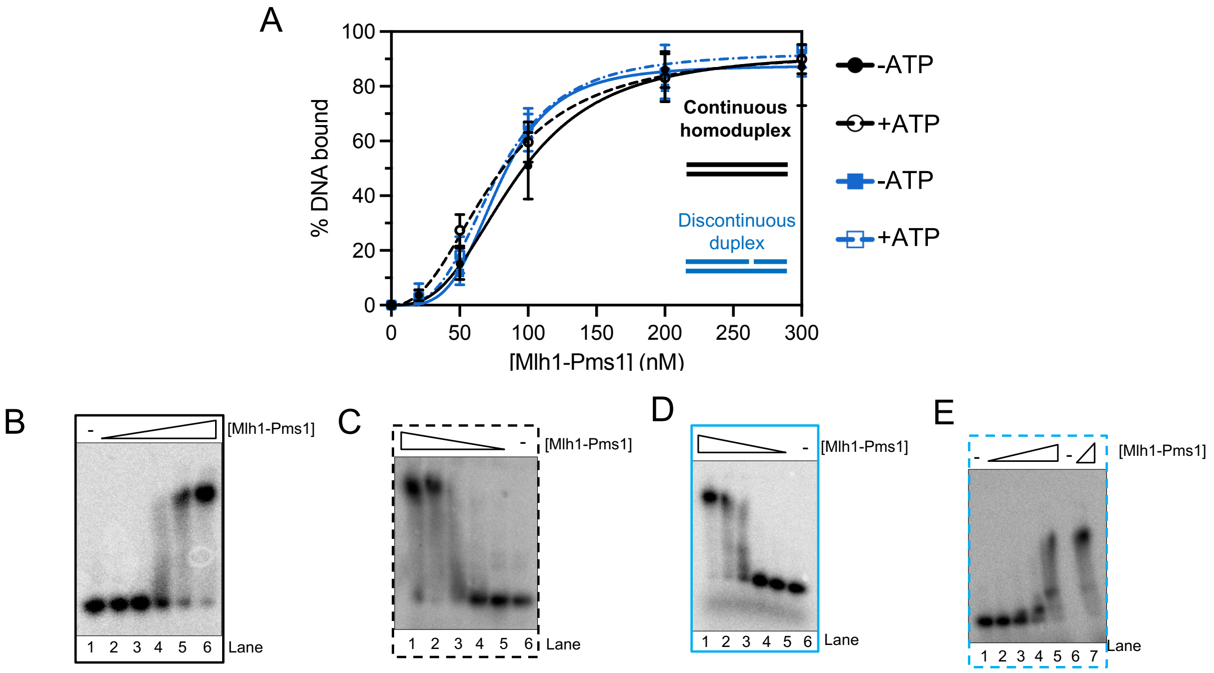


**Figure S2. Mlh1-Pms1 has modestly higher affinity for model oligonucleotide substrates containing a discontinuity than for intact duplex DNA.** (A) Quantification of electrophoretic mobility shift assays performed on intact (continuous) and discontinuous oligonucleotide substrates shown as percent DNA bound as described in the Materials and Methods. All reactions contained 0, 20, 50, 100, and 200 nM of Mlh1-Pms1 in 120 mM NaCl and 0.5 mM ATP when indicated. For all data collected, n = 3. Data were fit to a sigmoidal function describing cooperative ligand binding. (B-E) Representative native TBE gels to demonstrate the gel shift of Mlh1-Pms1 on radiolabeled oligonucleotide substrate mimicking intact DNA or DNA with a pre-existing discontinuity. Where indicated, ATP was added to a final concentration of 0.5 mM. (B) Mlh1-Pms1 affinity for intact (continuous) DNA duplex without ATP. Lanes 1-6 are in increasing order of concentration 0, 20, 50, 100, and 200 nM. (C) Mlh1-Pms1 affinity for intact (continuous) DNA duplexes with 0.5 mM ATP. Lanes 1-6 are in order of decreasing concentrations of 200, 100, 50, 20, and 0 nM Mlh1-Pms1. (D) Mlh1-Pms1 affinity for discontinuous duplexes without ATP. Lanes 1-6 are in order of decreasing concentrations of 200, 100, 50, 20, and 0 nM Mlh1-Pms1. (E) Mlh1-Pms1 affinity for discontinuous DNA duplexes with 0.5 mM ATP. Lanes 1-5 are in increasing concentration order from 0-200 nM Mlh1-Pms1, skipping lane 6, and 300 mM Mlh1-Pms1 in lane 7. Gels were quantified using ImageJ software. Due to unstable binding at near physiological ionic strength, the amount of DNA bound was quantified as loss of DNA in the substrate band relative to negative controls.

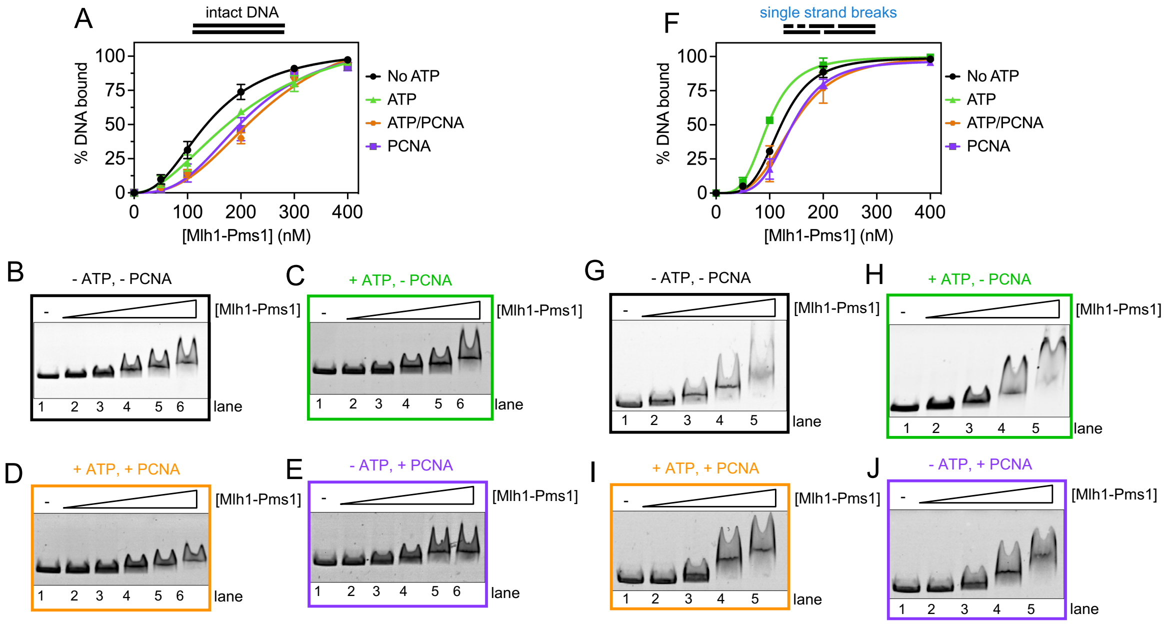


**Figure S3. Mlh1-Pms1 has higher affinity for linearized plasmid with single strand breaks relative to intact DNA.** (A) Quantification of electrophoretic mobility shift assays. The concentration of Mlh1-Pms1 used was 0 nM, 50 nM, 100 nM, 200 nM, and 400 nM for each experiment. Intact DNA was generated by linearizing pBR322 with HindIII. Data were fit to a sigmoidal function describing cooperative ligand binding. $K_d$ values are shown in Table 1. (B-E) Representative gels for DNA binding assay. (F) Same as A for linearized pBR322 with four single strand breaks generated using Nt.BstNBI. (G-H) Representative gels for DNA binding assay.

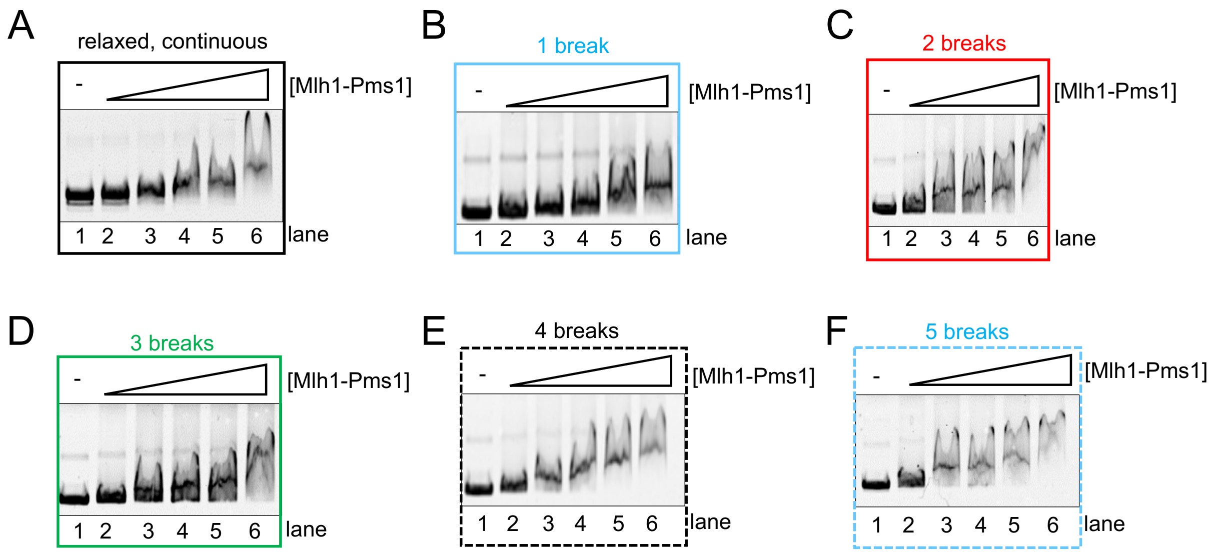


**Figure S4. Mlh1-Pms1's affinity for DNA increases as the number of single strand breaks increases.** Representative gels for experiments in Figure 1. Mlh1-Pms1 was titrated to final concentrations: 0, 50, 100, 150, 200 or 300 nM.

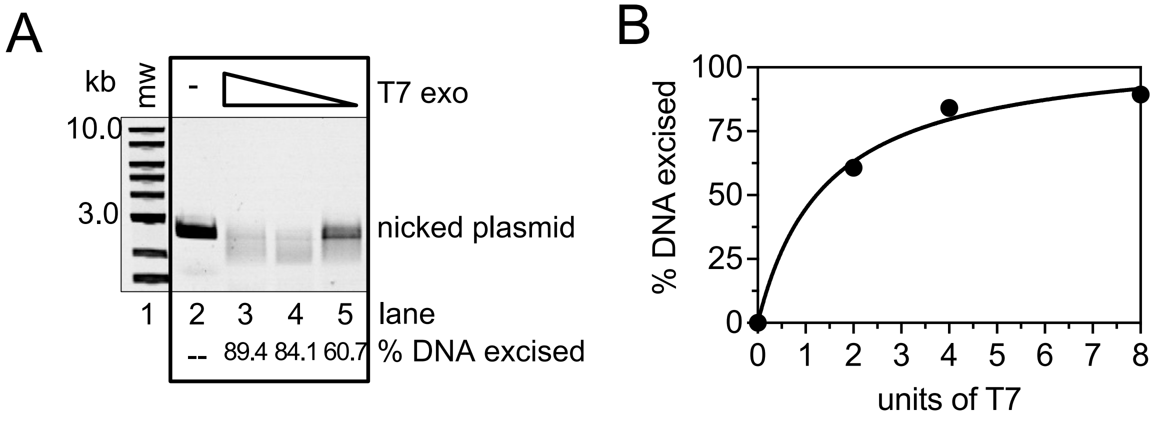


**Figure S5. Optimization of T7 exonuclease activity for Mlh1-Pms1 nick protection assays.** (A) T7 exonuclease was titrated in lanes 3-5 (2.0, 4.0, and 8.0 units) on a 2.7 kb DNA with a single pre-existing nick and analyzed by a native agarose gel to identify an amount of T7 that that partially degrades nicked DNA in the absence of other factors. Top band is intact double stranded circular DNA while lower band in gel is single stranded circular DNA strand after excision. (B) Quantification of data in A fit to a hyperbolic function. 2.0 units of T7 exonuclease was used for protection experiments in Figure 5.

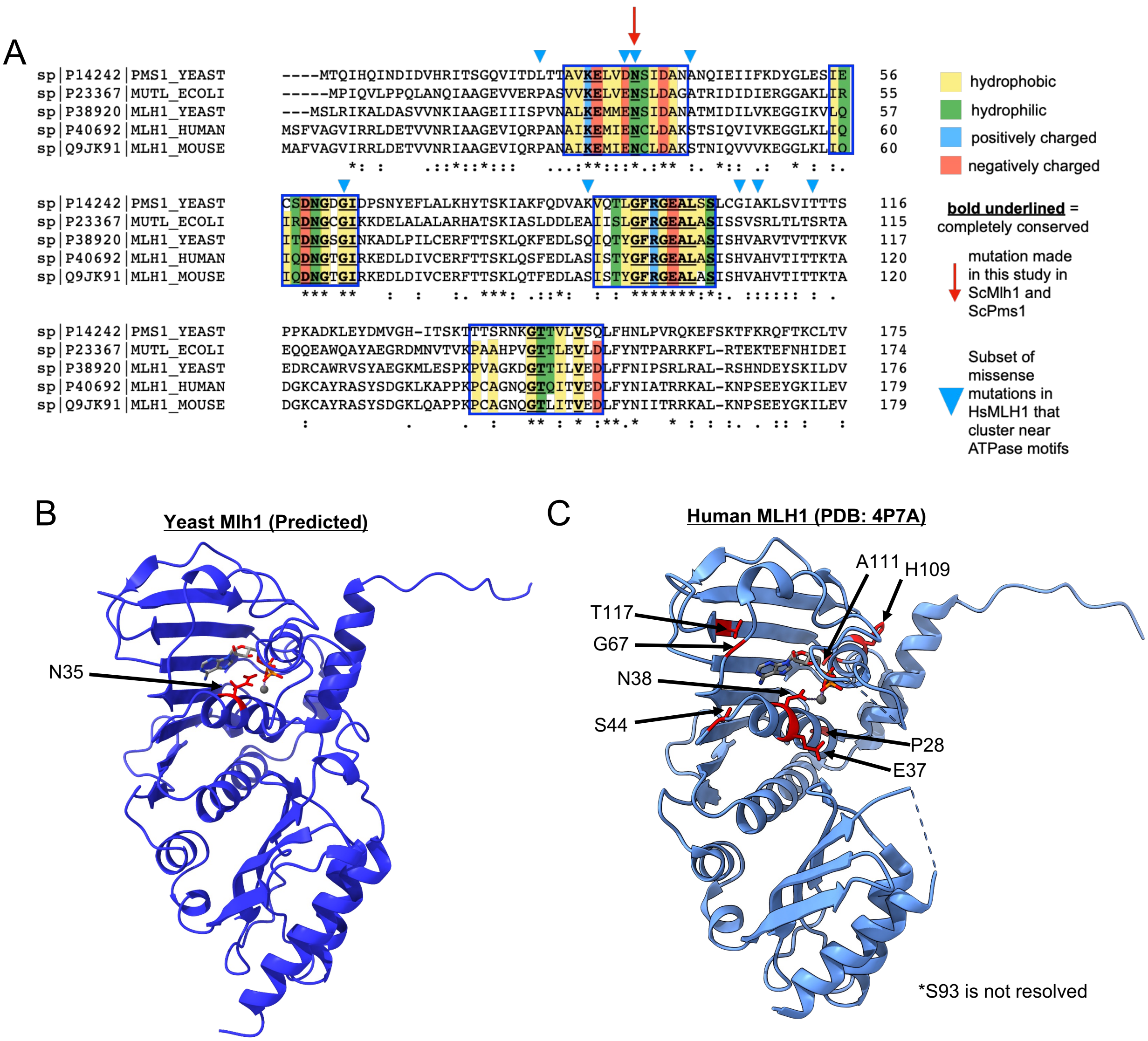


**Figure S6. The ATPase sites in Mlh1-Pms1 are conserved and biologically relevant.** (A) Clustal Omega Multiple Sequence Alignment of the first 179 amino acids from yeast Mlh1, yeast Pms1, *E. coli* MutL, human MLH1, and mouse MLH1. Boxed residues are the four ATPase motifs that are conserved among the GyrB, Hsp90, histidine kinase, MutL (GHKL) family (23, 26, 48) of ATPases. Residues within these motifs are color-coded and labeled for chemical properties to further demonstrate conservation. Red arrow indicates the position of the asparagine residue that was mutated in yeast Mlh1 and yeast Pms1 in this study to abolish ATPase activity. Blue triangles indicate a subset of missense mutations that have been identified in Lynch syndrome patients (43, 44) that cluster around ATPase motifs, including the conserved asparagine that is critical for ATPase activity. (B) Predicted structure of the amino-terminal domain of yeast Mlh1. AlphaFold2 was used predict the structure of the amino-terminal domain of yeast Mlh1. This was then aligned to crystal structure of the amino-terminal domain of *E. coli* MutL complexed with ADPNP (PDB: 1B63) (26) and to the crystal structure of the amino-terminal domain of human MLH1 complexed with ADP

(PDB: 4P7A) (49). Asparagine 35 in the predicted yeast structure aligned with the corresponding asparagine residues in the *E. coli* and human structures where it coordinates a metal ion. The magnesium ion and the ADP from the human structure along with the predicted yeast structure and the role of the N35 residue are shown here. (C) Crystal structure of the amino-terminal domain of human MLH1 complexed with magnesium and ADP. Missense mutations in panel A clustering around the ATPase motifs are mapped to the structure. These include N38 which corresponds to N35 in the yeast sequence and structure in B.
